# Nitric oxide signalling underlies salicylate-induced increases in neuronal firing in the inferior colliculus: a central mechanism of tinnitus?

**DOI:** 10.1101/2021.05.14.444151

**Authors:** Bas MJ Olthof, Dominika Lyzwa, Sarah E Gartside, Adrian Rees

## Abstract

The tinnitus-inducing agent salicylate reduces cochlear output but causes hyperactivity in higher auditory centres, including the inferior colliculus (the auditory midbrain). Using multi-electrode recording in anaesthetised guinea pigs (*Cavia porcellus*), we addressed the hypothesis that salicylate-induced hyperactivity in the inferior colliculus involves nitric oxide signalling secondary to increased ascending excitatory input.

In the inferior colliculus, systemic salicylate (200 mg/kg i.p., 0 h) markedly increased spontaneous and sound-driven neuronal firing (3-6 h post drug) with both onset and sustained responses to pure tones being massively increased. Reverse microdialysis of increasing concentrations of salicylate directly into the inferior colliculus (100 µM-10 mM, from 0 h) failed to mimic systemic salicylate. In contrast, it caused a small, transient, increase in sound-driven firing (1 h), followed by a larger sustained decrease in both spontaneous and sound-driven firing (2-5 h). When salicylate was given systemically, reverse microdialysis of the neuronal nitric oxide synthase inhibitor L-methyl arginine into the inferior colliculus (500 mM, 2-6 h) completely blocked the salicylate-induced increase in spontaneous and sound-driven neuronal firing.

Our data indicate that systemic salicylate induces neuronal hyperactivity in the auditory midbrain via a mechanism outside the inferior colliculus, presumably upstream in the auditory pathway; and that the mechanism is ultimately dependent on nitric oxide signalling within the inferior colliculus.

Given that nitric oxide is known to mediate NMDA receptor signalling in the inferior colliculus, we propose that salicylate activates an ascending glutamatergic input to the inferior colliculus and that this is an important mechanism underlying salicylate-induced tinnitus.

**Graphical abstract:** Tinnitus is associated with increased activity in the central auditory pathway, an effect replicated by high dose sodium salicylate. Here we show that salicylate-induced increases in neuronal activity in the auditory midbrain are mediated by nitric oxide signalling within this region. Nitric oxide is a key mediator of neuronal responses to NMDA receptor stimulation in this region. Our data support the hypothesis that tinnitus is mediated by increased ascending glutamatergic input to the auditory midbrain.

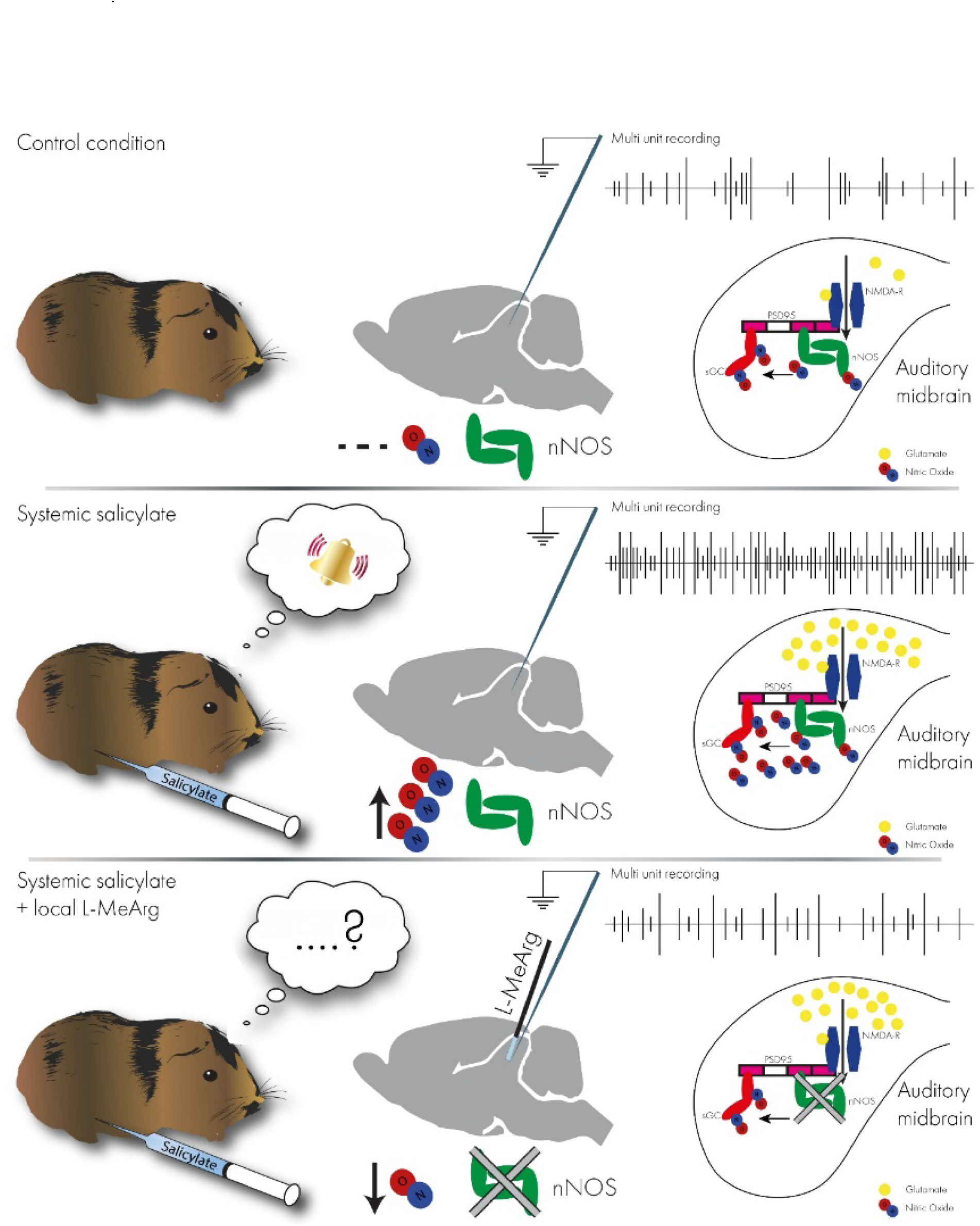

## 1. Introduction

Tinnitus, the sensation of a phantom sound, is experienced chronically by 10-15% of the population (Eggermont, 2012; Baguley *et al*., 2013). Although the onset of tinnitus is usually associated with some degree of peripheral hearing loss, the percept can persist after thresholds have returned to normal. One hypothesised mechanism underlying tinnitus is an enhancement in central gain. Thus, it is suggested that, in compensation for reduced peripheral input, central plasticity modifies the balance of excitation and inhibition at different levels in the auditory pathway leading to increased neural responsivity (Auerbach *et al*., 2014);(Norena, 2011);(Schaette & Kempter, 2006).

Tinnitus can occur in response to a variety of insults including exposure to loud sound, cochlear trauma, and therapeutic drugs. Salicylate, a non-steroidal anti-inflammatory drug, has long been known to induce tinnitus and hearing loss in humans, especially when taken in high doses (Mongan *et al*., 1973; Cazals, 2000). Systemic administration of salicylate has proven to be a valuable research tool in experimental animals because it allows the reliable and reversible induction of tinnitus possibly accompanied by hyperacusis (Jastreboff & Sasaki, 1986; Jastreboff *et al*., 1988; Jastreboff & Sasaki, 1994; von der Behrens, 2014). A primary effect of salicylate is to reduce cochlear output and thus induce hearing loss (Cazals, 2000). But salicylate induced tinnitus is also associated with a complex pattern of changes in spontaneous and sound driven firing at various levels in the ascending auditory system, including in the brainstem nuclei, the auditory midbrain, and the auditory cortex (Jastreboff & Sasaki, 1986; Chen & Jastreboff, 1995; Ochi & Eggermont, 1996; Eggermont & Kenmochi, 1998; Ma *et al*., 2006; Sun *et al*., 2009; Stolzberg *et al*., 2011; Chen *et al*., 2015; Jiang *et al*., 2017). However, questions remain as to what extent such changes are driven by the upstream consequences of salicylate’s effects on the cochlea or whether they represent direct effects of salicylate on neuronal activity in the central nervous system. For example, in the IC evidence points to multiple effects of salicylate in neural tissue including upregulation of NMDA receptor activity and down regulation of GABA and serotonin receptors (Bauer *et al*., 2000; Liu *et al*., 2003; Hwang *et al*., 2011; Hu *et al*., 2014).

Interestingly, accumulating evidence suggests that salicylate, and indeed other insults known to induce tinnitus, influence the expression of nNOS in the auditory pathway. Upregulation of nNOS in the ventral cochlear nucleus has been reported in animals in which tinnitus was confirmed behaviourally after either salicylate treatment (Zheng *et al*., 2006) or noise exposure (Coomber *et al*., 2014; Coomber *et al*., 2015). Furthermore, evidence for nitric oxide-dependent effects on firing in ventral cochlear nucleus neurons have been reported in animals with tinnitus (Hockley *et al*., 2020a). At the level of the auditory midbrain, we have reported increased nNOS expression in the dorsal cortex of the IC in rats with tinnitus (Maxwell *et al*., 2019).

The synthesis of nitric oxide by nNOS is known to be a step in the pathway that mediates glutamatergic excitatory signalling via NMDA receptors, and this mechanism widely implicated in neural plasticity (Garthwaite, 2008; Hardingham *et al*., 2013). We recently reported that most neurones in the central nucleus of the inferior colliculus (ICc) contain puncta of neuronal nitric oxide synthase (nNOS), and that these puncta are structurally associated with NMDA receptors (Olthof *et al*., 2019). Moreover, we provided evidence that glutamate signalling in the ICc, mediated via these NMDA receptors, depends on intact nitric oxide signalling. The role of these mechanisms in the physiology and pathophysiology of hearing remains unknown, but the upregulation of nNOS in tinnitus described above and evidence that NMDA receptor mRNA is also upregulated in the IC following noise trauma suggests that this signalling pathway may play a role in tinnitus (Dong *et al*., 2010a).

Here we examined the hypothesis that the increase in firing seen acutely in the IC after systemic salicylate administration is the result of increased nitric oxide signalling in the ICc secondary to enhanced glutamatergic stimulation of NMDA receptors. We defined the changes in neuronal firing in the ICc after systemic administration of salicylate in the guinea pig and delineated their time-course. We determined whether changes are mediated by salicylate acting within the IC or depend on up-stream pathways, and we examined whether they are dependent on the activation of the NMDA/nitric oxide signalling pathway in the ICc.

## 2. Methods

### 2.1 Animals

Male (n=2) and female (n=8) adult tricolour guinea pigs (*Cavia porcellus*) were bred in-house and housed in groups of 3-6 in spacious open pen housing under a 12 h light/dark cycle with controlled humidity. Animals were allowed dried food and water *ad libitum*, and their diet was supplemented by daily fresh fruit and vegetables. Animals were used as adults (42 - 92 days old).

### 2.2 Surgical procedure

Animals were anaesthetised with a combination of urethane (1 g/kg^-1^ i.p., Sigma), fentanyl (0.3 mg.kg^- 1 -^i.p., Hameln) and midazolam (5mg.kg^-1^ i.m., Hameln). Animals were monitored at regular intervals and received supplementary doses of fentanyl as required to prevent a withdrawal response to a hind toe pinch. Atropine sulphate (0.05 mg.kg^-1^ s.c.) was also administered before surgery to suppress bronchial secretions. A tracheotomy was performed, and the animal was allowed to respire freely in an atmosphere enriched with medical oxygen or was artificially respired as required with oxygen using a modified small animal respirator (Harvard Apparatus). The animal was transferred to a sound attenuated room and fixed in a modified stereotaxic frame (Kopf) equipped with hollow ear bars with Perspex specula positioned to allow an unobstructed view of the tympanic membrane. Body temperature was monitored with a rectal probe and was maintained at 38° C with a thermostatically controlled electric heating pad (Harvard Apparatus).

A craniotomy was made over the right IC, and the overlying cerebral cortex was aspirated. A concentric microdialysis probe manufactured in-house, see (Olthof *et al*., 2019) was implanted into the IC at an angle of 10° medio-lateral from the vertical, to a depth of approximately 4 mm below the surface of the IC. A 32-channel recording electrode with contact points arranged linearly over 3.2 mm (100 µm pitch; Neuronexus A1×32-10mm-100-177-A32), was implanted vertically in the IC immediately rostral to the dialysis probe. The depth of the electrode was adjusted to capture responses to the widest range of sound frequencies based on frequency response areas (Palmer *et al*., 2013) derived from the recorded firing. The microdialysis probe was constantly perfused (2 μl/min) with artificial cerebrospinal fluid (aCSF) (composition (mM): NaCl 140, KCl 3, 184 Na_2_HPO_4_ 0.707, NaH_2_PO_4_ 0.272, MgCl_2_ 1, D-glucose 10, and CaCl_2_ 2.4).

### 2.3 Sound stimulation and electrophysiological recording

The electrode was connected to a head stage (NN32AC Tucker Davis Technologies (TDT)) connected to a 32-channel preamplifier (PZ2-32, TDT), which in turn was connected by an optical interface to a BioAmp (RZ2, TDT). Electrophysiological data were sampled at 24.414 kHz. Multi-unit firing was recorded from all 32 channels in response to pure tones (75 ms duration, 10 ms rise/fall time, total sweep duration 150 ms). Tones were generated by a Multi I/O Processor (RZ6; TDT) at a sampling rate of 97.656 kHz and delivered to the animal via Sony MDR 464 earphones housed in alloy enclosures coupled to damped probe tubes that fitted into the Perspex specula (Rees, 1990). The maximum output of the system was approximately flat from 0.1 to 9 kHz (100 ± 8 dB SPL) above which it declined with a slope of ∼20 dB/octave such that the maximum output of the system at 16 kHz was 80 dB SPL. MATLAB™ scripts were used to control stimulus presentation and data recording, and to store recorded data for offline analysis.

#### Frequency response areas

(FRA) were generated from the multiunit firing recorded in response to the random presentation of pure tones varying between 256 Hz and 16.384 kHz in 1/10th octave intervals and at attenuations between 15 and 95 dB of the maximum output in 5dB steps. Each combination of frequency and level was presented once.

#### Peristimulus time histograms

(PSTHs) were generated from the multiunit firing recorded in response to the random presentation of 75-ms pure tones varying between 512 Hz and 8.192 kHz in octave intervals (hereafter referred to as 0.5, 1, 2, 4, and 8 kHz) and presented at 30 and 50 dB of attenuation (approximately 70- and 50-dB SPL respectively) with 150 sweeps recorded for each combination. Randomly interspersed in this sequence, 150 sweeps at maximal attenuation (120 dB, silent sweeps) were recorded to capture spontaneous firing.

### 2.4 Drug application

Following implantation, the microdialysis probe was perfused with aCSF for at least an hour to allow for stabilisation of the preparation. Beginning after this stabilization period (defined as 0 h), electrophysiological recordings were made for up to six hours. In experiment 1, the ‘systemic salicylate only’ condition (n=4), animals received a bolus dose of salicylate (200 mg.kg^-1^ i.p. at 4 ml.kg^-1^ in 0.1M PBS at 0 h); in this condition the microdialysis probe was perfused with aCSF throughout the experiment. In experiment 2, the ‘local salicylate’ condition (n=3), the microdialysis probe was perfused with 100 μM salicylate from 0-2 h, 1 mM salicylate from 2-4 h and 10 mM salicylate from 4-5h. In experiment 3 the ‘systemic salicylate & local L-MeArg’ condition (n=3), animals received a bolus injection of salicylate at 0 h and from 2-4 h the microdialysis probe was perfused with L-methyl arginine (Cambridge Bioscience; L-MeArg, 500 µM in aCSF) - a concentration previously shown to effectively inhibit nNOS *in vivo* (Garthwaite *et al*., 1989; Olthof *et al*., 2019).

### 2.5 Data processing

Electrophysiological recordings were bandpass filtered (300 Hz to 3000 Hz) and thresholded at 3 times the standard deviation of the spontaneous firing during the baseline block collected immediately prior to pharmacological interventions. Firing was defined as the spikes exceeding this threshold. Only electrode sites which showed clear sound-driven firing, and which represented the tonotopic sequence in the central nucleus of the IC, were included in the data analysis (14-19 electrode sites per animal).

To delineate the excitatory region of the FRA (eFRA), the number of spikes per 150 ms sweep elicited by a single presentation was determined for each combination of frequency and level. For display purposes, each square in a grid (representing tone frequencies on the X-axis and the loudness levels on the Y-axis) was colour coded according to a fixed colour scale to indicate the firing level. To determine the eFRA, we derived an outline based on 3 x the standard deviation of the average firing during maximally attenuated (‘silent’) responses on that electrode site. This was smoothed with a Gaussian filter and processed to include only contiguous blocks of at least 25 grid squares. This outline was used to derive the threshold, characteristic frequency (CF), and eFRA width (at threshold +30 dB).

PSTHs were constructed from 150 sweeps with a bin width of 1 ms and a duration of 150 ms. To measure spontaneous firing, the average number of spikes per sweep during 150 silent sweeps was calculated. For PSTHs, the average number of spikes per sweep was calculated for 150 presentations of the stimuli at each of five frequencies and two levels (totalling 10 frequency/level combinations). ‘Total’ firing refers to the total number of spikes per sweep in the whole 150 ms sweep. ‘Onset’ firing refers to the number of spikes per sweep in first 20 ms after the onset of the tone-driven response (determined for each electrode site), and ‘sustained’ firing is that in the final 20 ms. ‘Post-stimulus’ firing is the number of spikes per sweep between 80-150 ms (i.e. from 5 ms after the end of the stimulus to the end of the sweep). As recorded firing could vary greatly between electrode sites, we also present the change in firing detected at each electrode site as a ratio of that detected at baseline (0 h).

### 2.6 Experimental Design and Statistical Analysis

The data sets in experiments 1, 2, and 3 comprised responses recorded at 72 electrode sites (n=4 animals), 48 electrode sites (n=3 animals) and 49 electrode sites (n=3 animals), respectively. To examine the time-course of the effect of pharmacological interventions on spontaneous and sound-driven firing, the response at each hourly time point post drug administration was compared to the firing at baseline (0 h) using the non-parametric Friedman one-way repeated measures ANOVA followed by post hoc Wilcoxon signed rank tests. The p values from the Friedman ANOVA were corrected for the number of stimulus conditions tested (10 + spontaneous firing, n=11) in each experiment. The p values from the Wilcoxon test were corrected for multiple comparisons within each frequency/level combination (n=6 or 5, hourly time points compared to baseline). Corrections were performed using the Holm-Šídák method. In addition, we compared the systemic salicylate alone and systemic salicylate & local L-MeArg conditions. Spontaneous firing at each time point was compared using a two-way mixed model ANOVA (with ‘Treatment’ as a between groups factor and ‘Time’ as a within groups factor). Sound-driven firing was compared using a four-way mixed model ANOVA (with ‘Treatment’ as a between groups factor and ‘Time’, ‘Frequency’ and ‘Attenuation’ as within groups factors). Significant (Treatment x Time) interaction terms were followed up with planned comparisons between groups at each time point by t-tests (Bonferroni corrected). For non-spherical data (Mauchley’s test), degrees of freedom were corrected by the Greenhouse-Geisser method. All statistical analyses were performed using IBM SPSS Statistics version 25.

## 3. Results

We were able to identify electrode sites lying within the ICc (as distinct from the ICD or ICL) by the presence of a tonotopic distribution of multi-unit firing in the eFRAs generated at each electrode site. Data from between 14 and 19 electrode sites per animal in the ICc were included in the analysis. The eFRAs recorded with multi-channel silicone electrodes were relatively homogenous and V shaped. This contrasts with the variety of eFRA shapes for single units recorded with tungsten or glass electrodes we and others have previously reported (LeBeau *et al*., 2001; Palmer *et al*., 2013; Orton & Rees, 2014; Vogler *et al*., 2014). The difference most likely reflects the fact that the relatively low impedance silicone electrodes, used here, pick up multiple units such that an average V shaped eFRA dominates, whereas higher impedance electrodes can pick up spikes from single units which vary in the shape of their eFRA.

### 3.1 Experiment 1: Systemic administration of salicylate enhances spontaneous and sound-driven firing in the ICc

Following the systemic injection of salicylate, we observed an increase in firing in the ICc (Fig 1). In the FRA, increases were seen within the eFRA (Figure 1 white outline) which equates to well-tuned sound-driven firing, as well as outside the eFRA. This effect on firing developed over several hours following salicylate administration. Marked increases in firing were evident at 3 hours post salicylate and by 6 hours, firing was massively increased. Although the eFRA on some of the electrode sites shifted slightly to higher frequencies and/or lower thresholds, this was inconsistent between electrode sites, and overall, there was no change in eFRA parameters (threshold, CF, or bandwidth). From the FRA alone, it is not possible to determine whether increased firing (particularly that outside the eFRA) is spontaneous firing, unrelated to sound stimuli, or represents a reduction in frequency selectivity. Moreover, it is not possible to see when within the 150-ms sound-driven response changes occurred. To investigate the nature of the response to salicylate further, we derived PSTHs to measure spontaneous firing in silent sweeps and sound-driven firing in response to five selected frequencies and two sound levels lying within and outside the eFRA (Figure 1 blue markers).

**Figure 1.**
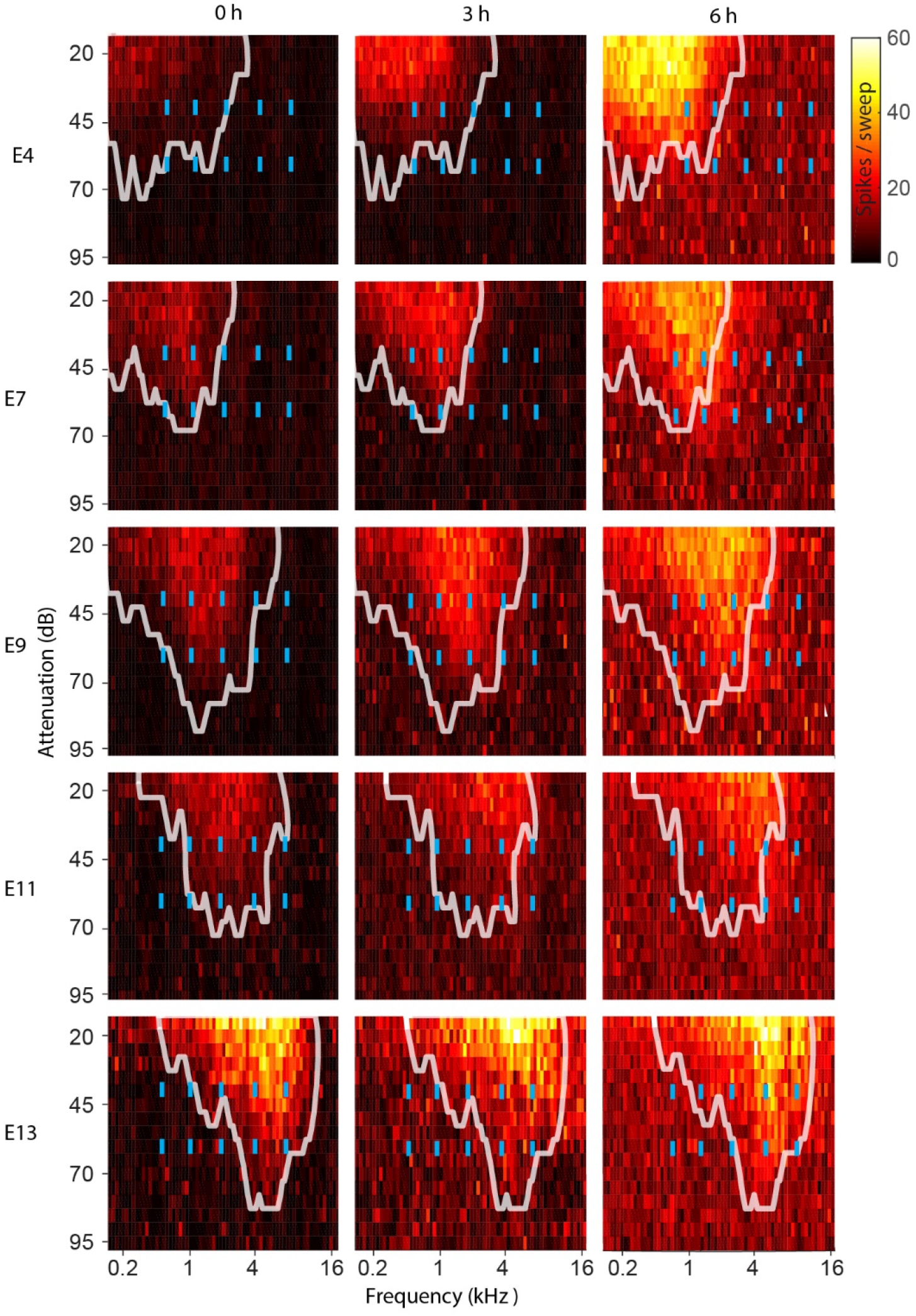
Systemic salicylate administration increases firing observed in frequency response area (FRA) plots in the ICc. Example FRA plots from five electrode sites at increasing depth from the dorsal surface within the ICc(E4, E7, E9, E11, E13) in one animal. The figure shows the dorsal to ventral tonotopic representation of sound-driven firing in the ICc, and the impact of systemic salicylate at 3 h and 6 h relative to baseline (0 h). The white outline denotes the eFRA at baseline. Note that, following administration of salicylate, there is an increase in firing (see key) within the eFRA (outline) as well as at attenuations and frequencies outside the eFRA. Blue markers indicate location of PSTH stimuli in the FRA space (see later).

#### Spontaneous firing

Based on the PSTHs generated in response to silent stimuli, we found that after salicylate injection there was a small decrease in spontaneous firing at 1 and 2 hours (Fig 2b and Table 1) followed by a progressive increase up to the end of the recording at 6 hours (Fig 2a-c and Fig 2d). These changes were seen across the ICc at deep as well as at more superficial recording locations. Whereas at baseline, spontaneous firing was always below 2 spikes/sweep (Fig 2ai), 6 hours after salicylate administration, spontaneous firing was as high as 5 spikes/sweep (Fig 2aiii). As baseline spontaneous firing varied between electrode sites (partly as a function of likely differences in impedance of the electrode sites and potentially as a function of the electrode site’s proximity to active units), we also show the change in firing detected at each electrode site as a ratio of that at baseline (Fig 2b). Statistical analysis of the raw data (Friedman ANOVA followed by post-hoc Wilcoxon signed ranks compared to baseline) showed that spontaneous firing was slightly, but significantly, decreased 1 h and 2 h after systemic salicylate administration, but was significantly increased by 3 h and increased further over the next 3 h remaining significantly above baseline up to the end of the experiment at 6 h (Table 1).

**Table 1.**
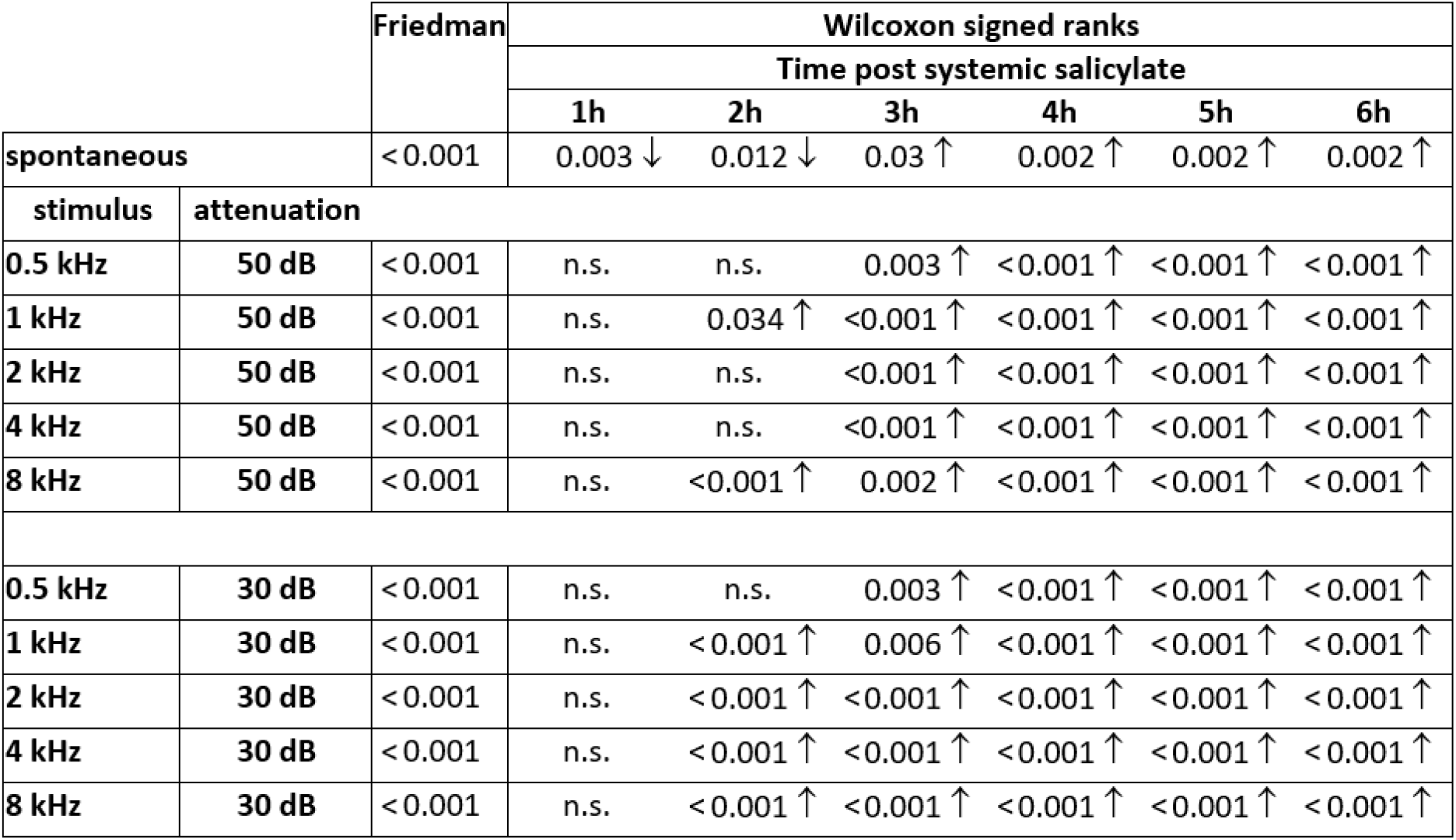
Experiment 1. Total firing in the PSTH following systemic salicylate: Friedman ANOVA and Wilcoxon signed rank post hoc test statistics. Friedman one-way ANOVA for each frequency and attenuation, p values were corrected for the eleven frequency/attenuation comparisons. Significant ANOVA was followed by post hoc Wilcoxon signed ranks comparisons between baseline and each post-salicylate time point for each frequency at each attenuation, p values were corrected for these six comparisons. Arrows indicate direction of change.

**Figure 2.**
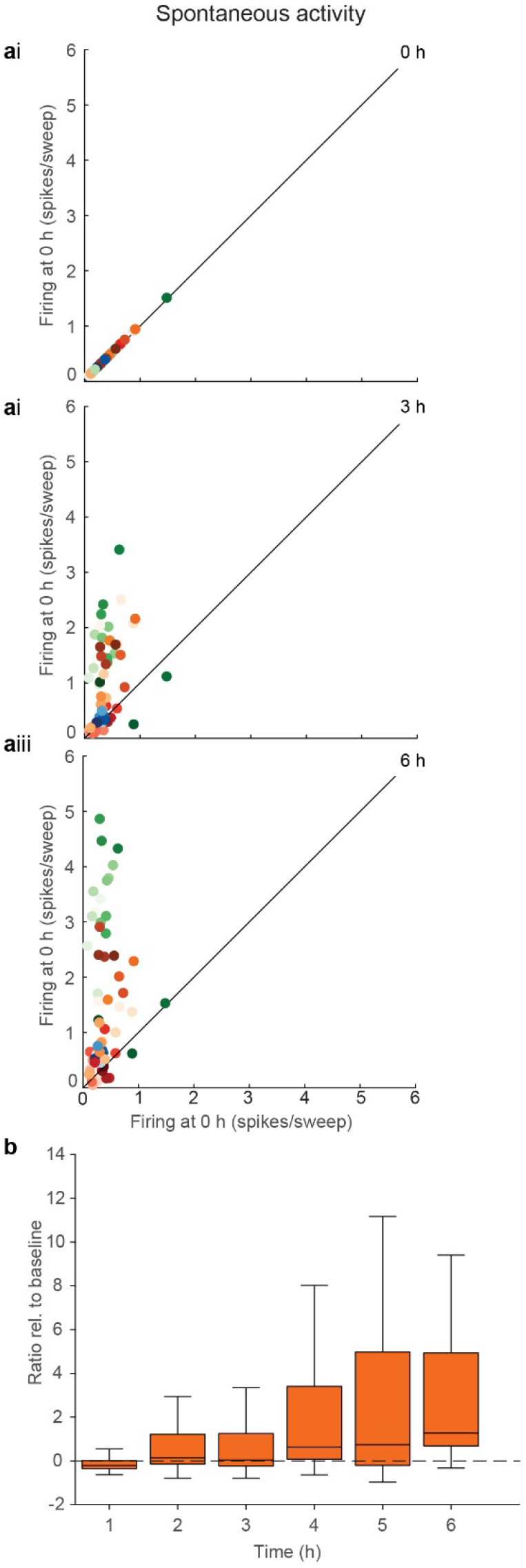
Experiment 1. Systemic salicylate administration evokes a persistent increase in spontaneous firing in the ICc. a) Correlation of spontaneous firing at i) baseline (0 h), ii) 3 h, and iii) 6 h with baseline firing at 0 h. Green, red, orange, and blue symbols represent different animals; dark colours are data from more dorsally located electrode sites, lighter colours are data from more ventrally located electrode sites. As time after administration progresses, more points are found above the line of unity indicating an increase in firing relative to baseline. Note: data from many of the electrode sites overlie each other in these plots. Data at each time point represent the average spikes/sweep for 150, 150-ms sweeps. b) Group data showing spontaneous firing rates from 1-6 h following systemic administration of salicylate. Box plots show median and interquartile range (box) and maximum and minimum observations (whiskers). Changes in firing are normalised to firing at baseline. Data are from 72 electrode sites in 4 animals.

#### Sound-driven firing

In addition to the changes in spontaneous firing, systemic salicylate also evoked an increase in sound-driven firing across the ICc(Figure 3ai-iii, 3b and 3c). This first became evident for some frequencies and attenuations at 2 h and was apparent for all frequencies and attenuations at 3h. Sound-driven firing remained elevated up to 6 h (Figure 3ai-iii, 3b and 3c).

**Figure 3.**
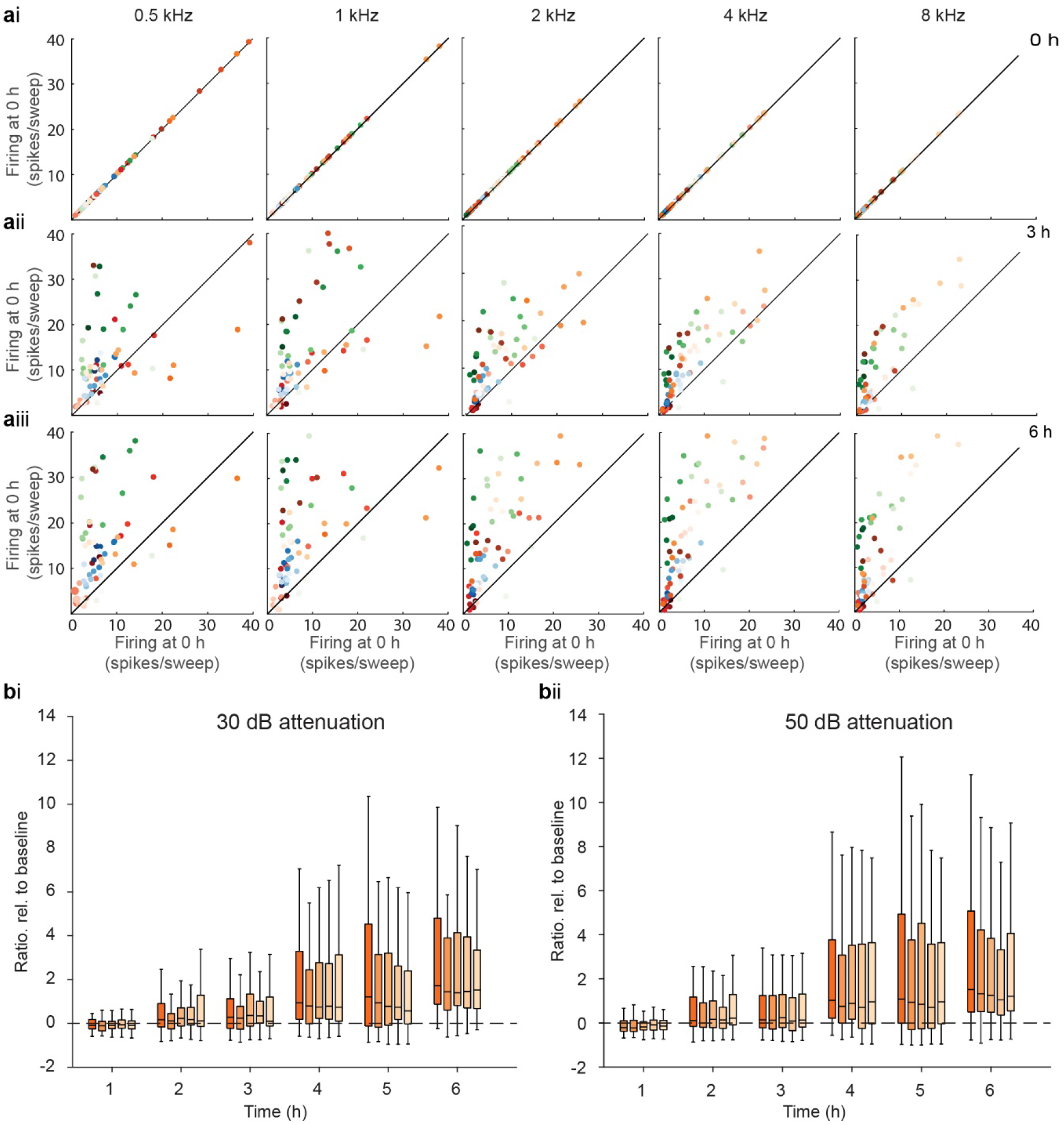
Experiment 1. Systemic salicylate administration evokes a persistent increase in sound-driven firing in the ICc. Total firing in the PSTH in response to stimuli at five frequencies following systemic administration of salicylate. a) Correlation of total firing at i) baseline (0 h), ii) 3 h, and iii) 6 h with firing at 0 h for stimuli at 30 dB attenuation. Green, red, orange, and blue symbols represent different animals; dark colours represent data from electrode sites located dorsally and lighter colours represent data from electrode sites located more ventrally. As time after administration progresses, more points are found above the line of unity indicating an increase in firing relative to baseline. Data at each time point represent the average spikes/sweep over 150, 150-ms sweeps. b) Grouped data showing total firing in the PSTHs in response to pure tone stimuli at five frequencies at i) 30 dB and ii) 50 dB attenuation measured hourly across the 6h period following systemic administration of salicylate. Box plots show median and interquartile range (box) and maximum and minimum observations (whiskers). Changes in firing are normalised to firing at baseline. Data are from 72 electrode sites in 4 animals.

Statistical analysis of the total firing revealed that, for all frequencies and both attenuations, there was a highly significant increase in firing over time (Table 1). Post-hoc Wilcoxon signed ranks revealed that for most frequencies at 30 dB attenuation firing was significantly increased 2 h post injection and remained significantly elevated up to 6 h. When stimuli were presented at a lower sound level (50 dB attenuation), similar effects were seen.

The data in Figure 3bi-ii and each row in Table 1 is based on the response of all electrode sites to the specified stimulus. However, owing to the marked tonotopic organisation of the ICc, the response to a particular frequency is stronger on one electrode than on others, as is evident in Figure 4 (central diagonal). When we examined the effect of salicylate on the responses to individual frequencies measured at specific electrode sites, we observed that firing was enhanced for tones at the BF for that electrode site (among the five frequencies presented). However, it was notable that the greatest effect of salicylate was often at a frequency above or below the BF at that sound level (Figure 4, yellow shaded boxes). To quantify this effect, we compared the ratio of firing at baseline and after salicylate (6 h) for the BF (at baseline), to the ratios one frequency above and one frequency below BF. In both cases the increase in firing either side of BF was greater than at BF (median ratio at BF-1 vs BF= 2.32 vs 1.83, n= 48, p < 0.001 and median ratio at BF+1 vs at BF= 3.05 vs 1.89, n= 46 p < 0.001).

**Figure 4.**
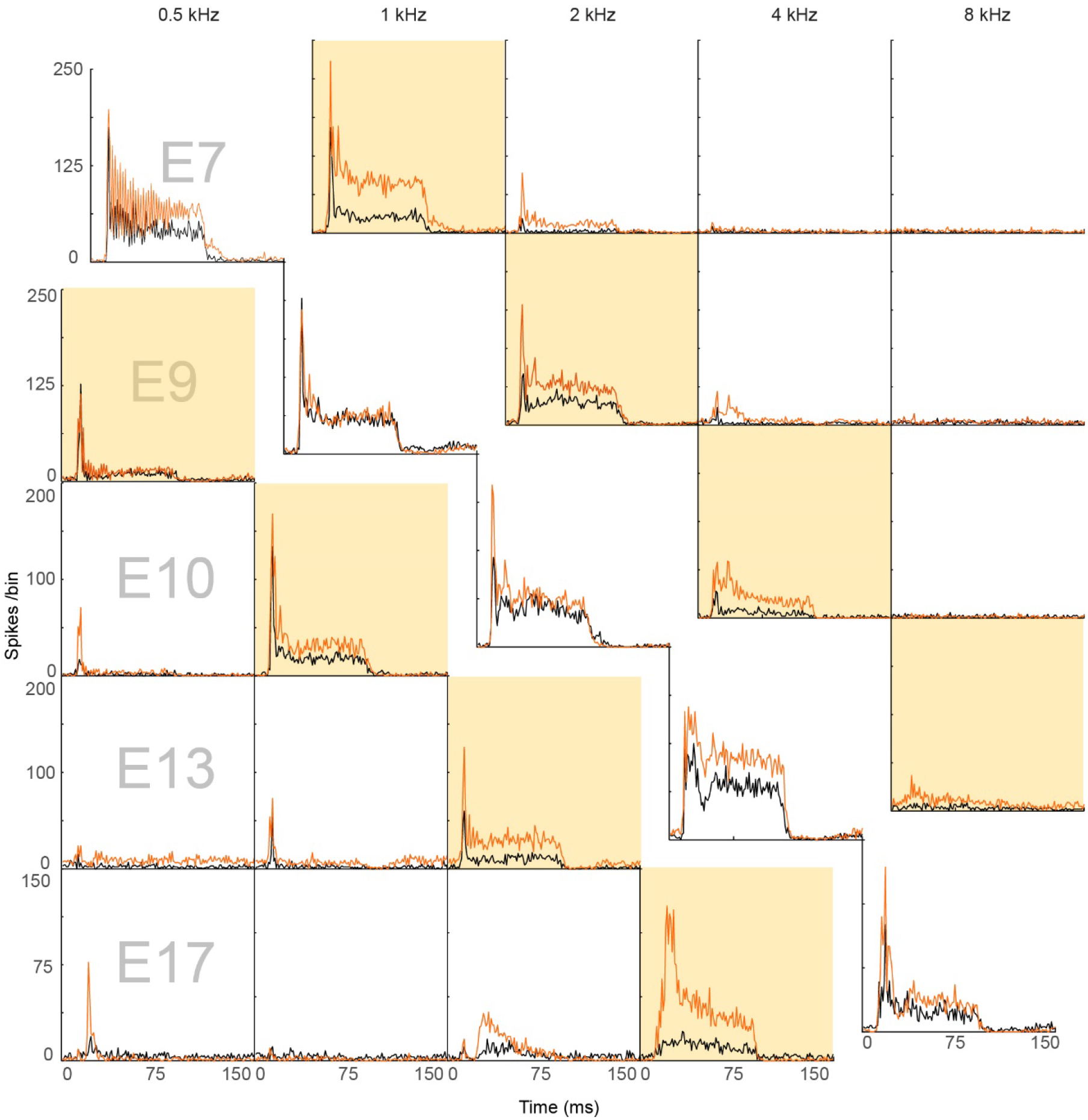
Experiment 1. Systemic salicylate increases sound-driven firing in the ICc at stimuli surrounding the best frequency. Example PSTHs from one animal showing responses to pure tone stimuli. Each row shows data from one electrode site (chosen as having the highest firing in response to one of the five stimulus frequencies). Each column represents the response to one frequency. Black lines are data collected at baseline (immediately before administration of systemic salicylate); orange lines are data collected 6 h after systemic salicylate. Note that following salicylate, the number of spikes in the PSTH is increased both for BF tones (i.e. those evoking the highest response for that electrode site), and tones which are lower and/or higher than the BF. Yellow boxes highlight responses to frequencies flanking the BF response of that electrode site at baseline.

As shown in Figure 4, following systemic salicylate the increase in firing occurred across both the onset and the sustained parts of the driven response with both showing a very similar magnitude effect to that for the total. Statistical analysis showed a similar pattern to that seen for the total firing (Table 1) with only a few differences in the times at which the increases in firing for particular stimulus parameters reached statistical significance (data not shown). An increase in firing was also seen in the low-level firing during the post-stimulus period which reflects post-stimulus effects and, at later times, spontaneous firing, although the actual numbers of spikes in this period was very low.

### 3.2 Experiment 2: Salicylate perfused locally into the IC does not mimic the effects of systemically administered salicylate

To determine whether any of the changes in firing in response to systemic administration of salicylate are mediated directly within the ICc, we examined the effect of salicylate perfused locally via a microdialysis probe at progressively increasing concentrations (0.1 mM 0-2 h; 1 mM 2-4 h; 10 mM 4-5 h).

#### Spontaneous firing

In contrast to the effect of systemically administered salicylate, salicylate perfused locally induced a small and transient increase in spontaneous firing in the first hour, followed by a more long-lasting reduction in firing which did not appear to be concentration dependent (Figure 5 ai-iii and b). Statistical analysis of the spontaneous firing showed a highly significant effect of local salicylate over time with a significant increase from baseline at 1 h and significant reductions at 2-5 h (Table 2).

**Table 2.** Experiment 2. Total firing in the PSTH following local perfusion of salicylate: Friedman ANOVA and Wilcoxon signed rank post hoc test statistics. Friedman one-way ANOVA for each frequency and attenuation: p values were corrected for the eleven frequency/attenuation comparisons. Significant ANOVA was followed by post hoc Wilcoxon signed ranks comparisons between baseline and each time point following initiation of local perfusion of salicylate for each frequency at each attenuation: p values were corrected for these six comparisons. Arrows indicate direction of change.

|  |  | Friedman | Wilcoxon signed ranks |  |  |  |  |
| --- | --- | --- | --- | --- | --- | --- | --- |
|  |  |  | Time (concentration) post local salicylate |  |  |  |  |
|  |  |  | 1h<br>(0.1 mM) | 2h<br>(0.1 mM) | 3h<br>(1 mM) | 4h<br>(1 mM) | 5h<br>(10 mM) |
| spontaneous |  | <0.001 | 0.004↑ | 0.004↓ | 0.004↓ | 0.016↓ | 0.016↓ |
| stimulus | attenuation |  |  |  |  |  |  |
| 0.5 kHz | 50 dB | <0.001 | 0.001↑ | <0.001↓ | <0.001↓ | 0.010↓ | 0.010↓ |
| 1 kHz | 50 dB | <0.001 | <0.001↑ | 0.006↓ | <0.001↓ | 0.007↓ | 0.006↓ |
| 2 kHz | 50 dB | <0.001 | <0.001↑ | 0.003↓ | <0.001↓ | 0.004↓ | 0.004↓ |
| 4 kHz | 50 dB | <0.001 | 0.008↑ | 0.008↓ | <0.001↓ | 0.008↓ | 0.008↓ |
| 8 kHz | 50 dB | <0.001 | 0.006↑ | 0.004↓ | <0.001↓ | 0.006↓ | 0.006↓ |
| 0.5 kHz | 30 dB | <0.001 | <0.001↑ | 0.018↓ | <0.001↓ | 0.018↓ | 0.018↓ |
| 1 kHz | 30 dB | <0.001 | 0.005↑ | 0.031↓ | 0.005↓ | 0.026↓ | 0.005↓ |
| 2 kHz | 30 dB | <0.001 | 0.005↑ | 0.008↓ | 0.005↓ | 0.008↓ | 0.006↓ |
| 4 kHz | 30 dB | <0.001 | 0.008↑ | 0.008↓ | 0.005↓ | 0.008↓ | 0.008↓ |
| 8 kHz | 30 dB | <0.001 | 0.008↑ | 0.008↓ | <0.001↓ | <0.001↓ | 0.006↓ |

**Figure 5.**
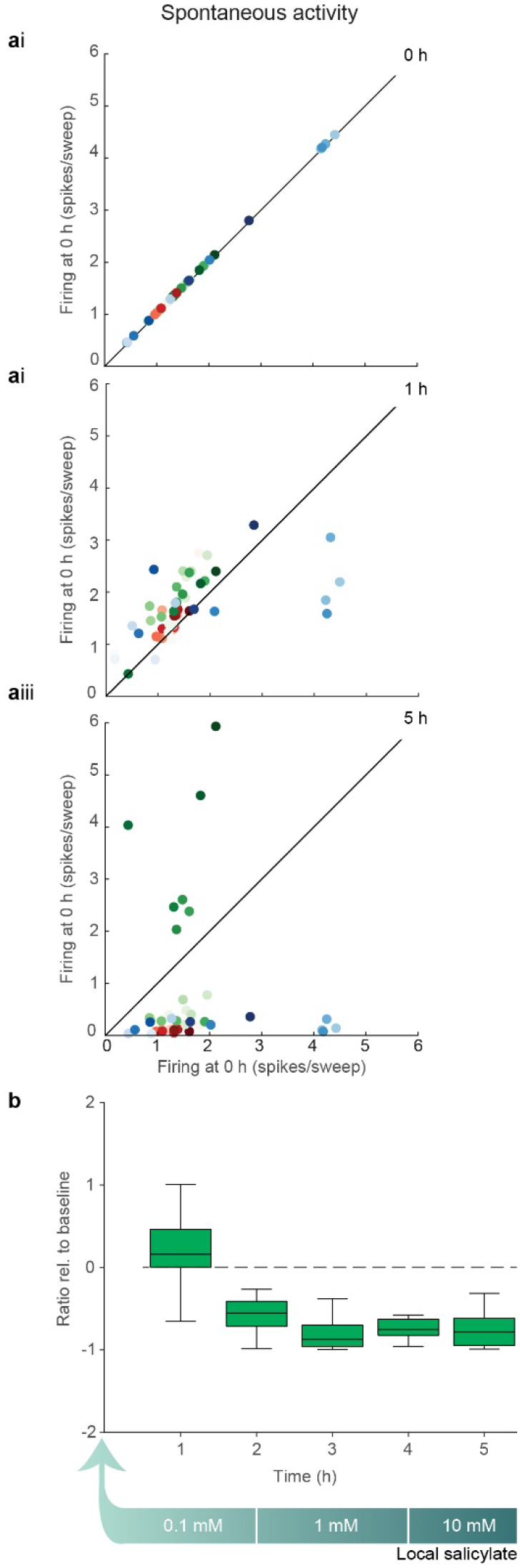
Experiment 2. Salicylate perfused locally into the IC causes a persistent decrease in spontaneous firing in the ICc. a) Correlation of spontaneous firing at i) baseline (0 h), ii) 1 h, and iii) 5 h with firing at 0 h. Green, red, and blue symbols represent 3 different animals; dark colours are data from more dorsally located electrode sites, lighter colours are data from more ventrally located electrode sites. At 1 h, most points are above the line of unity indicating increased firing whilst at 5 h, most points are below the line of unity indicating decreased firing. Data at each time point represent the average spikes/sweep for 150, 150-ms sweeps. b) Group data showing spontaneous firing rates from 1-5 h following local perfusion of salicylate at the concentrations shown (green arrow). Box plots show median and interquartile range (box) and maximum and minimum observations (whiskers). Changes in firing are normalised to firing at baseline. Data are from 49 electrode sites in 3 animals.

#### Sound-driven firing

Local salicylate evoked the same changes in sound-driven firing as in spontaneous firing (Figure 5). It was of note that during the first hour of perfusion of 0.1 mM salicylate there was a small increase in sound-driven firing, but during the second hour of perfusion of this concentration, as well as higher concentrations (1 and 10 mM), firing decreased (Figure 6 aii-iii, 6b and 6c). These effects were evident at both sound levels used. Both the transient increase and the long-lasting decrease in the firing following local salicylate were evident in both the onset and the sustained parts of the response (data not shown).

**Figure 6.**
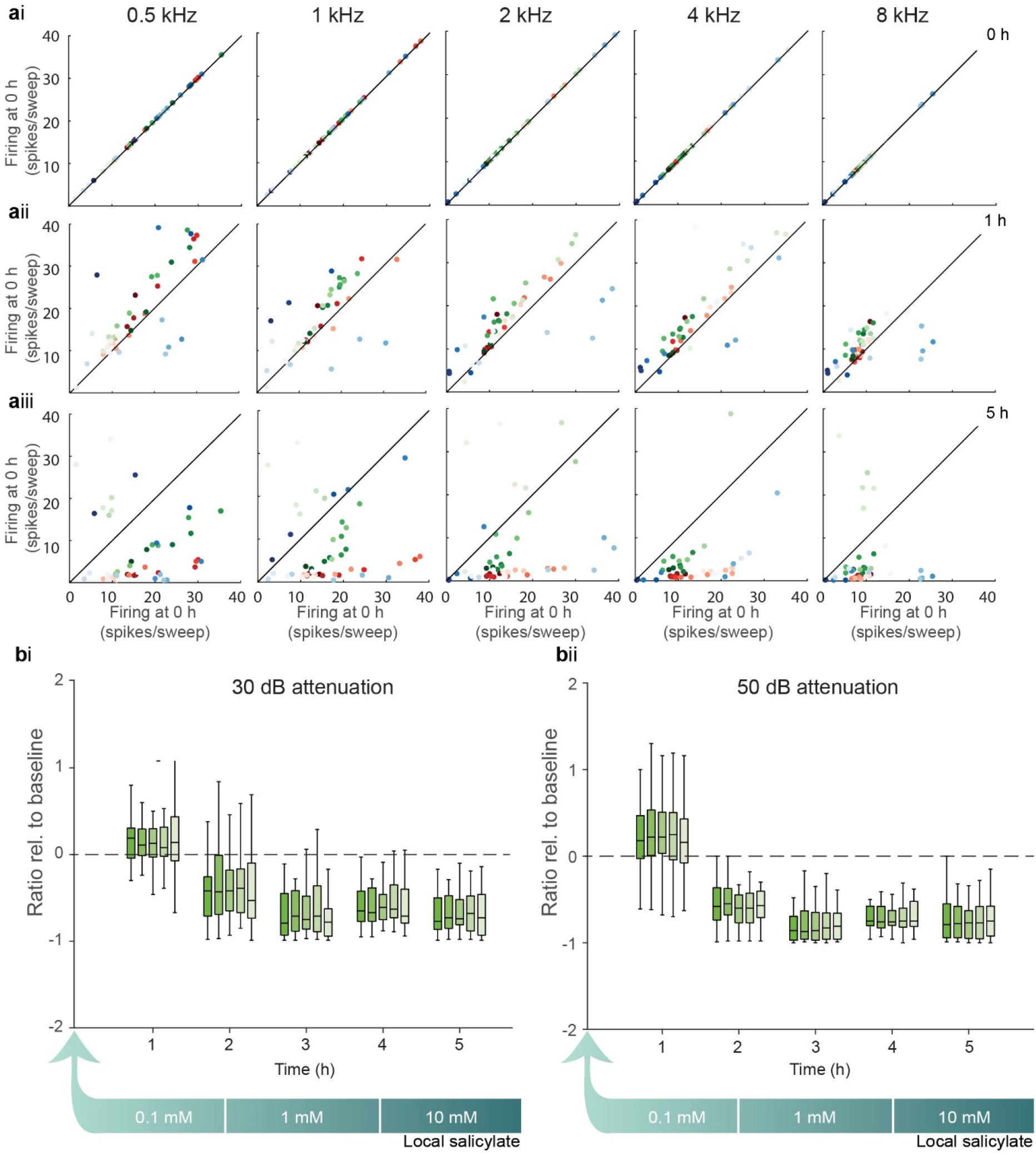
Experiment 2. Local perfusion of salicylate causes a persistent decrease in sound-driven firing in the ICc. Effect of locally perfused salicylate on total firing in the PSTH in response to pure tone stimuli at five frequencies. a) Correlation of firing at i) baseline (0 h), ii) 1 h, and iii) 5 h with firing at 0 h for stimuli at 30 dB attenuation. Green, red, and blue symbols represent 3 different animals; dark colours represent data from electrode sites located dorsally and lighter colours represent data from electrode sites located more ventrally. Note that at 1 h, most points are above the line of unity indicating increased firing whilst at 5 h, most points are below the line of unity indicating decreased firing. Data at each time point represent the average spikes/sweep for 150, 150-ms sweeps. b) Grouped data showing total firing in the PSTHs in response to pure tone stimuli at five frequencies at i) 30 dB and ii) 50 dB attenuation across the 5 h period during local perfusion of salicylate at the concentrations shown (green arrow). Box plots show median and interquartile range (box) and maximum and minimum (whiskers). Changes in firing are normalised to firing at baseline. Data are from 49 electrode sites in 3 animals.

Analysis of the total firing rates showed highly significant effects of local salicylate at all frequencies and both sound levels and post hoc tests revealed significant differences from baseline at all time points after initiation of local salicylate perfusion at all frequencies and both sound levels (Table 2).

### 3.3 Experiment 3: L-MeArg perfused locally into the IC blocks the effects of systemic salicylate

We hypothesised that the increase in spontaneous and sound-driven firing in the ICcin response to systemically administered salicylate is mediated by a mechanism involving nitric oxide signalling within the ICc. To test this, we perfused the nNOS inhibitor L-MeArg locally into the IC via a microdialysis probe beginning two hours after systemic injection of salicylate. Note, in Experiment 1 a microdialysis probe was also implanted but perfused with aCSF throughout the experiment.

FRAs comparing the effects of systemic salicylate alone with those of salicylate with the local perfusion of L-MeArg show that the increases in both spontaneous and sound-driven firing observed with systemic salicylate alone were abolished when L-MeArg was perfused locally into the IC (Figure 7).

**Figure 7.**
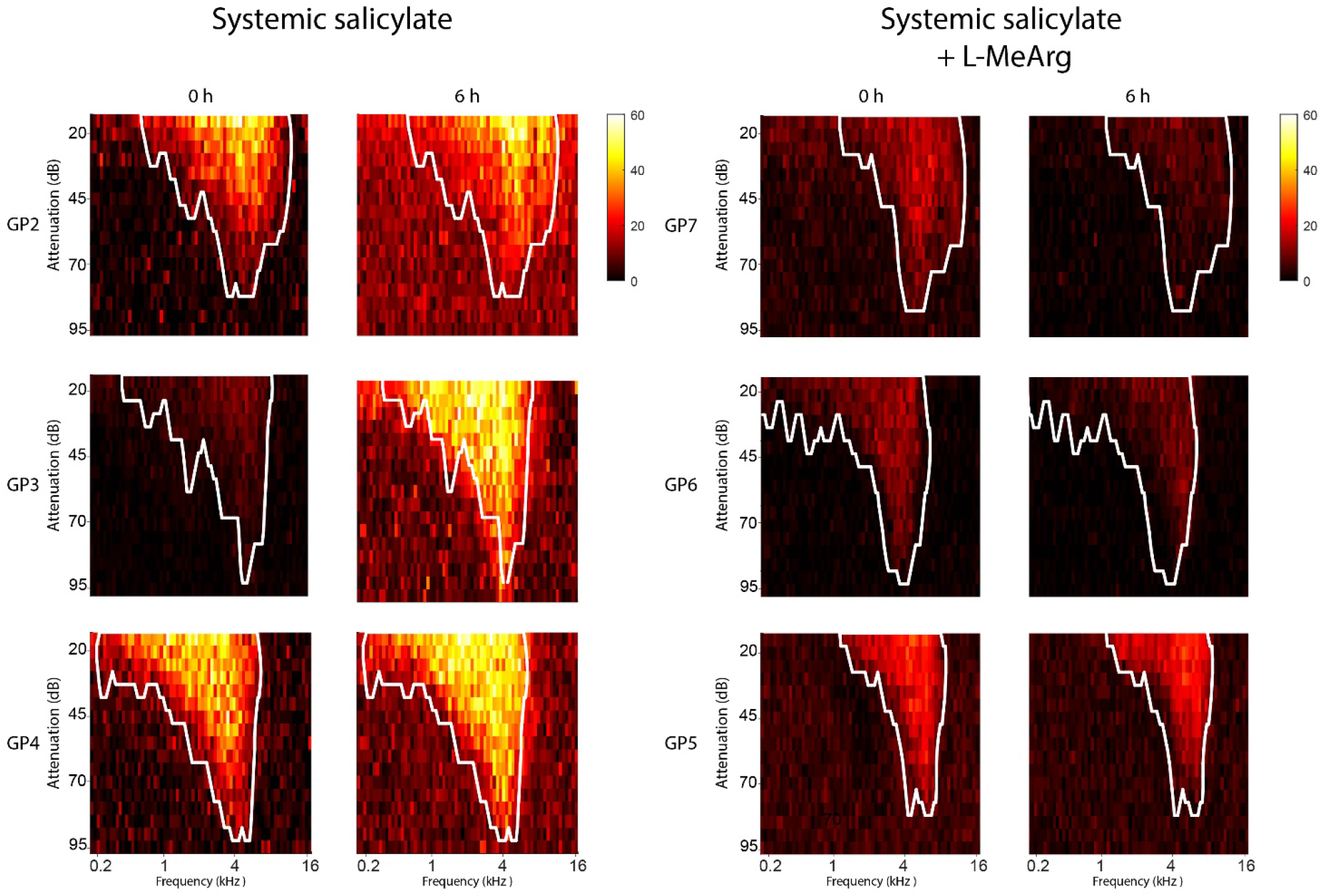
Experiment 1 and 3. The increased firing observed in the FRA following systemic salicylate administration is blocked by local perfusion of L-MeArg. Example FRA plots from a single electrode site in three animals from each of the two experimental groups approximately matched for best frequency. In each plot, the white outline denotes the eFRA at baseline. Note that, 6h following systemic administration of salicylate, there is an increase in driven firing within the eFRA (outline) and in spontaneous firing outside the eFRA. These effects of systemic salicylate do not occur when L-MeArg is perfused locally in the IC from 2-6 h.

#### Spontaneous firing

The local perfusion of L-MeArg blocked the increase in spontaneous firing seen with systemic salicylate alone. As shown in Figure 8ai-iii and 8b (scaled for comparison with Figure 2), when L-MeArg was perfused after the systemic administration of salicylate, the spontaneous firing did not increase relative to baseline. Although statistical analysis showed a significant effect over time, post hoc tests revealed that spontaneous firing differed only at the 3 h time point when it was actually slightly decreased compared to baseline (Table 3).

**Table 3.** Experiment 3. Total firing in the PSTH following systemic salicylate plus L-MeArg: Friedman ANOVA and Wilcoxon signed rank post hoc test statistics. Friedman one-way ANOVA for each frequency and attenuation, p values were corrected for the eleven frequency/attenuation comparisons. Significant ANOVA was followed by post hoc Wilcoxon signed ranks comparisons between baseline and each time point following administration of salicylate for each frequency at each attenuation; L- MeArg was perfused locally from 2-6 h. p values were Holm-Šídák corrected for the six comparisons. Arrows indicate direction of change.

|  |  | Friedman | Wilcoxon signed ranks |  |  |  |  |  |
| --- | --- | --- | --- | --- | --- | --- | --- | --- |
|  |  |  | Time post systemic salicylate (L-MeArg perfused from 2-6 h) |  |  |  |  |  |
|  |  |  | 1h | 2h | 3h | 4h | 5h | 6h |
| spontaneous |  | <0.001 | n.s. | n.s. | 0.000↓ | n.s. | n.s. | n.s. |
| stimulus | attenuation |  |  |  |  |  |  |  |
| 0.5 kHz | 50 dB | <0.001 | n.s. | n.s. | 0.006 ↓ | n.s. | n.s. | n.s. |
| 1 kHz | 50 dB | <0.001 | n.s. | n.s. | n.s. | n.s. | n.s. | n.s. |
| 2 kHz | 50 dB | <0.001 | n.s. | n.s. | 0.012 ↓ | n.s. | n.s. | n.s. |
| 4 kHz | 50 dB | <0.001 | n.s. | n.s. | 0.018 ↓ | n.s. | n.s. | n.s. |
| 8 kHz | 50 dB | <0.01 | n.s. | n.s. | n.s. | n.s. | n.s. | n.s. |
| 0.5 kHz | 30 dB | <0.001 | n.s. | n.s. | n.s. | n.s. | n.s. | n.s. |
| 1 kHz | 30 dB | <0.001 | n.s. | n.s. | n.s. | n.s. | n.s. | n.s. |
| 2 kHz | 30 dB | <0.01 | n.s. | 0.047 ↑ | n.s. | n.s. | n.s. | n.s. |
| 4 kHz | 30 dB | <0.001 | n.s. | n.s. | n.s. | n.s. | n.s. | n.s. |
| 8 kHz | 30 dB | <0.001 | n.s. | 0.012 ↑ | n.s. | n.s. | n.s. | n.s. |

**Figure 8.**
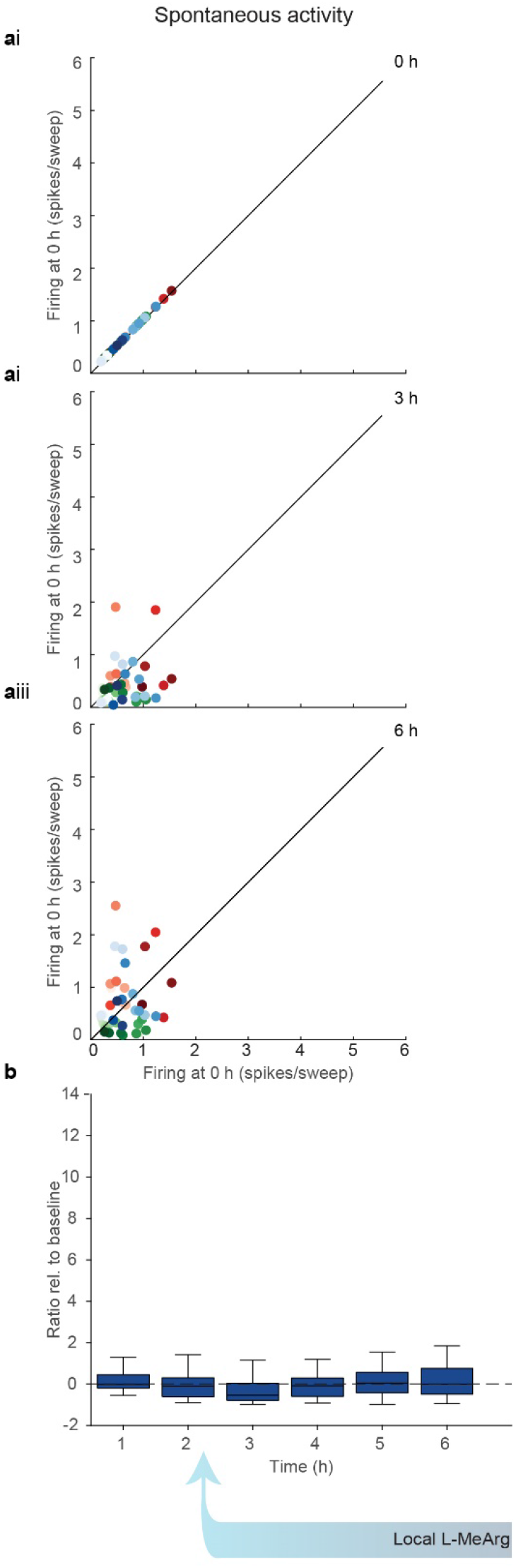
Experiment 3. Local perfusion of L-MeArg blocks the systemic salicylate-induced increase in spontaneous firing in the ICc. a) Correlation of spontaneous firing at i) baseline (0 h), ii) 3 h, and iii) 6 h post injection of salicylate with firing at 0 h; L-MeArg was perfused from 2-6 h. Green, red, and blue symbols represent different animals; dark colours are data from more dorsally located electrode sites, lighter colours are data from more ventrally located electrode sites. Data at each time point represent the average spikes/sweep for 150, 150-ms sweeps. b) Grouped data showing spontaneous firing rates from 1-6 h following systemic administration of salicylate; L-MeArg was perfused from 2-6 h. Box plots show median and interquartile range (box) and maximum and minimum observations (whiskers). Changes in firing are normalised to firing at baseline. The ordinate is scaled to that of Fig 3 to facilitate comparison. Data are from 48 electrode sites in 3 animals.

#### Sound-driven firing

Local perfusion of L-MeArg also blocked the increase in sound-driven firing induced by systemic salicylate (Figure 9). As shown in Figure 9iii (Scaled for comparison with Figure 3), during the period when L-MeArg was perfused (i.e. from 3 h after systemic salicylate), there were only minimal changes in sound-driven firing at 30 dB attenuation. For lower sound level stimuli, responses were actually decreased at some frequencies (Figure 9bii). Statistical analysis showed that there was a significant effect over time at all frequencies and attenuations. Post hoc tests showed that after initiation of local L-MeArg perfusion, sound driven firing did not change significantly from baseline in response to 30dB stimuli but showed a transient decrease at 3 h in response to some frequencies at 50 dB (Table 3).

**Figure 9.**
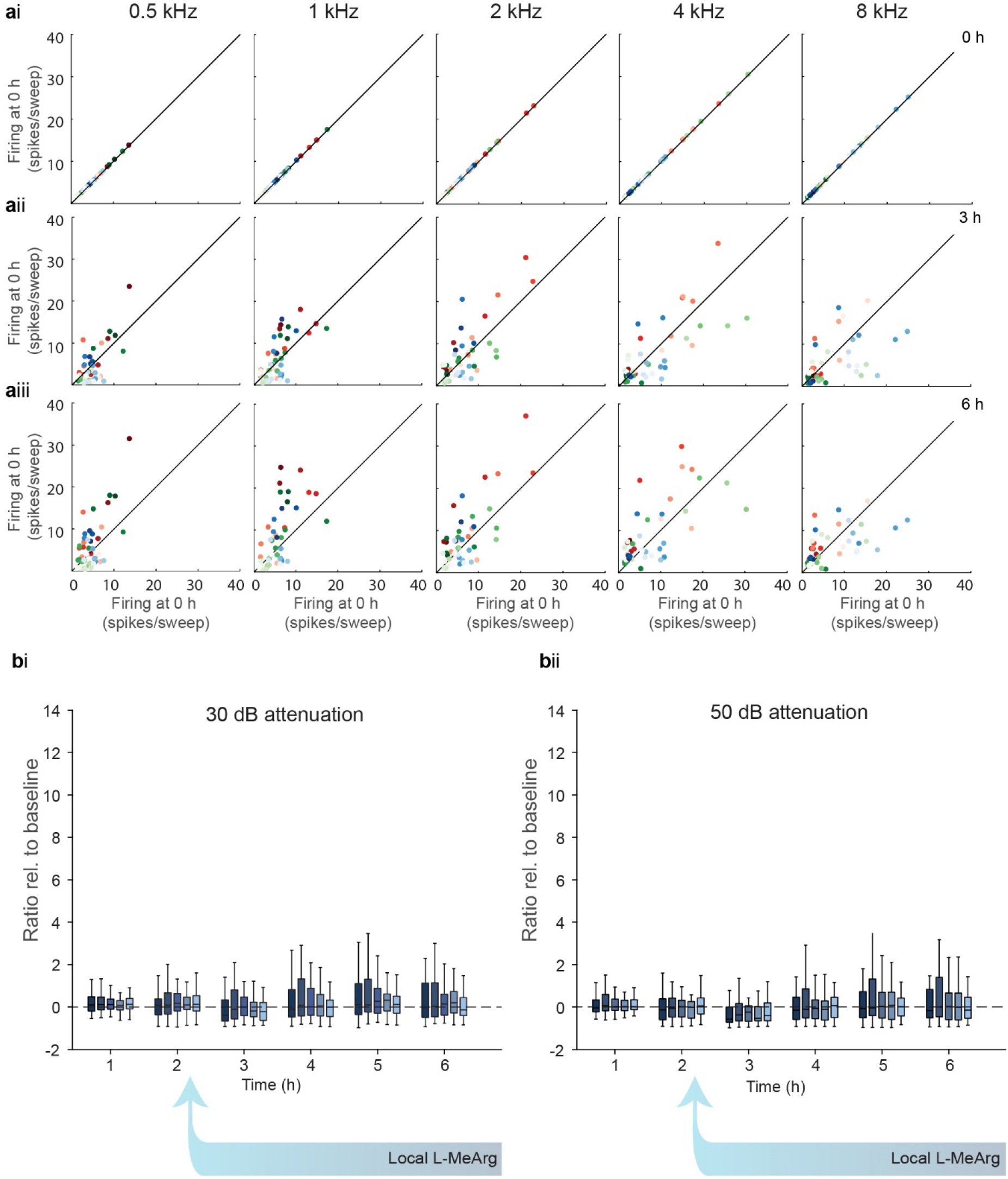
Local perfusion of L-MeArg blocks the systemic salicylate induced increase in sound-driven firing in the ICc. Effect of systemic salicylate plus local perfusion of L-MeArg on total firing in the PSTH in response to pure tone stimuli at five frequencies. a) Correlation of firing at i) baseline (0 h), ii) 3 h, and iii) 6 h with firing at 0 h for stimuli at 30 dB attenuation; L-MeArg was perfused from 2-6 h. Green, red, and blue symbols represent 3 different animals; dark colours represent data from electrode sites located dorsally and lighter colours represent data from electrode sites located more ventrally. Note that at 3h and 6 h points are scattered equally above and below the line of unity indicating no consistent change in firing relative to baseline. Data at each time point represent the average spikes/sweep for 150, 150-ms sweeps. b) Grouped data showing total firing in the PSTHs in response to pure tone stimuli at five frequencies at i) 30 dB and ii) 50 dB attenuation across the 6h period following systemic administration of salicylate; L-MeArg was perfused from 2-6 h. Box plots show median and interquartile range (box) and maximum and minimum observations (whiskers). Changes in firing are normalised to firing at baseline. The ordinate is scaled to that of Fig 3b to facilitate comparison. Data are from 48 electrode sites in 3 animals.

### 3.4 Between groups analysis: Experiment 1 and Experiment 3

We used between groups analysis to compare the systemic salicylate alone group (experiment 1) with the systemic salicylate plus local L-MeArg group (experiment 3) for both spontaneous and sound-driven firing (Figure 10a and b).

**Figure 10.**
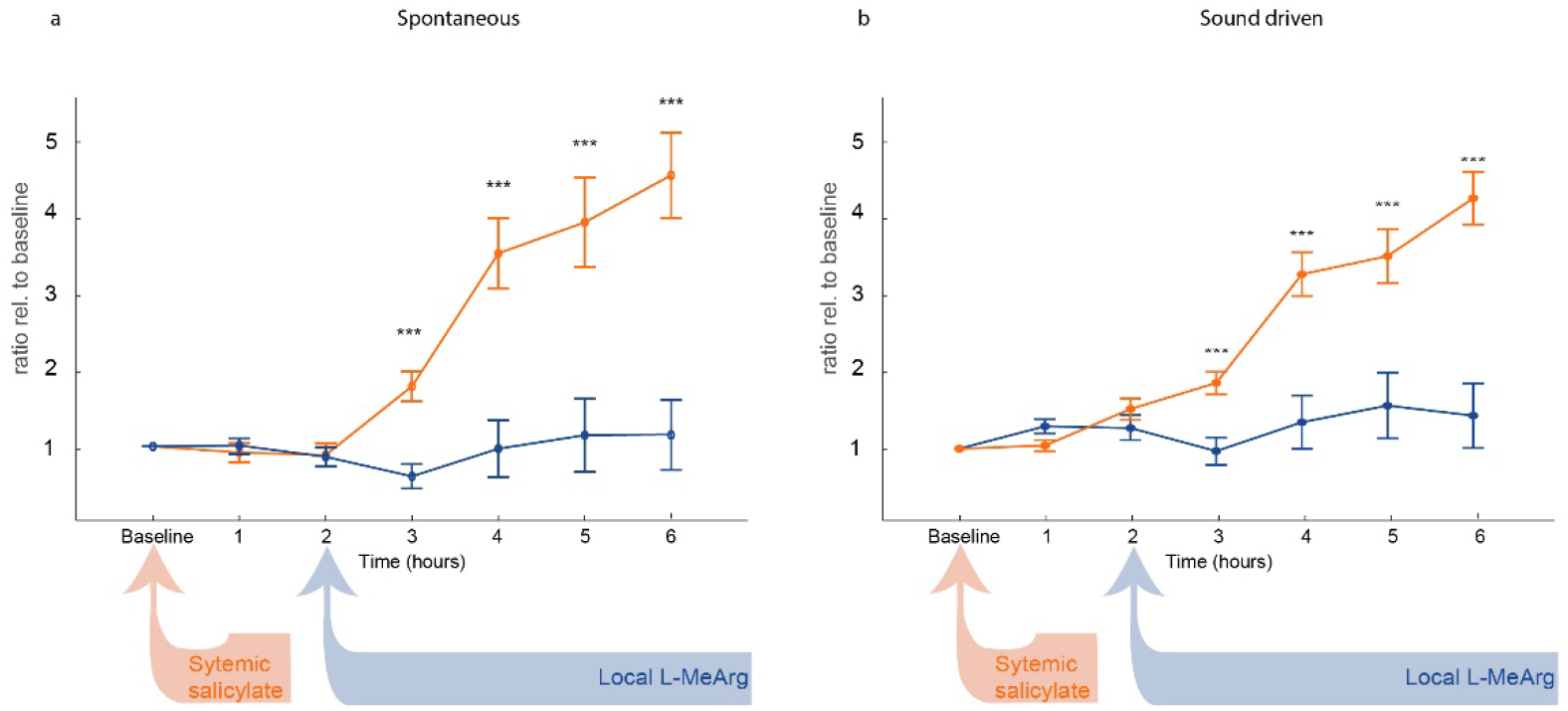
Experiment 1 vs experiment 3. L-MeArg locally perfused into the IC blocks the effect of systemic salicylate on spontaneous and sound-driven firing in the ICc. Comparison of the effect of systemic salicylate alone (orange) and systemic salicylate & local L-MeArg (blue) on a) spontaneous and b) sound-driven firing in the ICc. Data are mean ± sem average from 72 electrodes in 4 animals (orange) and 48 electrode sites in 3 animals (blue); for sound-driven data all frequencies/attenuations are averaged. Changes in firing are normalised to firing at baseline. ***p <0.001 Bonferroni corrected unpaired t test (assuming unequal variance) following significant ANOVA. See text for full statistical analysis.

#### Spontaneous firing

A two-way mixed model ANOVA with between groups factor ‘Treatment’ (salicylate alone vs salicylate + L-MeArg) and within groups factor ‘Time’ (hours post salicylate injection) showed significant main effects of Treatment (F_1,118_ = 23.45; p = 4 x 10^−6^) and Time (F_1.35,159.0_ = 24.09; p = 12 x 10^−9^) and a significant Treatment x Time interaction (F_1.35,159.0_ = 28.648, p = 9.33 x 10^−9^). Post hoc tests showed that the two groups were not different at 1 h or 2 h, but were significantly different at 3-6 h, i.e. during the period when L-MeArg was perfused (3 h: p = 7.2 x 10^−5^; 4 h: p = 0.00139; 5 h: p = 0.00254; 6 h: p = 1.8 x 10^−5^, Bonferroni corrected).

#### Sound-driven firing

A four-way mixed model ANOVA with a between group factor of ‘Treatment’, and within groups factors of ‘Time’, ‘Attenuation’ (30 dB vs 50 dB) and ‘Frequency’ (0.5-8 kHz) revealed a significant main effect of Treatment (F_1,118_ = 18.38, p = 3.7 x 10^−4^), together with significant main effects of Time (F_1.56, 183.89_ = 29.14, p = 6.7 x 10^−10^) and Attenuation (F_1,118_ = 7.85, p = 0.006). There was also a significant Time x Treatment interaction (F_1.56, 183.9_ = 18.39, p = 7.8 x 10^−7^). Post hoc t-tests indicated that the two treatments were not different at 1 and 2h, but differed significantly at 3, 4, 5, and 6 h (i.e. during L-MeArg perfusion) (3 h: p = 3.8 x 10^−4^; 4 h: p = 1.8 x 10^−5^; 5 h: p = 4.68 x 10^−4^; 6 h: p = 1.2 x 10^−7^, Bonferroni corrected).

## 4. Discussion

Here we tested the hypothesis that systemic salicylate increases neural firing in the auditory midbrain through a nitric oxide dependent mechanism. Systemic salicylate, at a high dose known to induce tinnitus (Jastreboff *et al*., 1988; Brozoski & Bauer, 2016), increased both spontaneous and sound-driven firing in the ICcfrom two hours until the end of the experiment at six hours. These effects were not mimicked when salicylate was delivered directly into the ICcvia a microdialysis probe, suggesting that they are mediated outside (presumably upstream) of the ICc. The effects of systemically administered salicylate on both spontaneous and sound-driven firing were completely prevented when we delivered the nNOS inhibitor L-MeArg directly into the ICc. These data indicate that within the ICcthe effects of systemic salicylate are mediated by nitric oxide signalling. Given that this dose of salicylate has been shown to cause tinnitus and hyperacusis in guinea pig, our data are consistent with the hypothesis that that nitric oxide signalling in the IC is an important mediator of these phenomena.

### 4.1 Systemic salicylate increases neuronal firing in the ICc two hours following administration

Previous studies describing increases in firing in the IC following salicylate have only reported data from either a single time point or unspecified times within a time window after salicylate (Jastreboff & Sasaki, 1986; Chen & Jastreboff, 1995; Ma *et al*., 2006; Sun *et al*., 2009). Here using electrodes that allow for stable multi-unit recording over a prolonged period, we extended those electrophysiological findings by tracking the changes in firing at specified times from the same recording sites over several hours after systemic salicylate administration. The increased firing rates we observed with salicylate began about 2 hours after administration and increased progressively over the 6 h of recording with no sign of plateauing up to that time. Spontaneous firing rates were reduced significantly below baseline levels for the first 2 hours before showing significant increases at later time points. Similarly, driven rates also only increased after the first 2 hours post salicylate. That there was no increase in the driven firing for 2 hours after the salicylate injection is consistent with Sun *et al*. (2009) who reported no change in the LFP evoked by tones in the IC in awake rat 1 hour after systemic salicylate. The large increase in sound-driven firing in the ICc evoked by systemic salicylate occurred across all frequencies and levels. It is important to note that the increase in driven firing does not simply reflect the increase in spontaneous firing as the absolute increase in driven firing greatly exceeded the change in spontaneous firing. Our multi-unit firing measure likely included units that respond to sound with different time courses. We cannot distinguish the responses of these different types, but systemic salicylate increased the sound-evoked firing in both the onset and sustained periods of the driven response. Although, this enhanced response most likely represents increased firing in those units that are active at baseline, it is possible that it also includes a contribution from previously silent units. Our findings of increased spontaneous and driven firing are consistent with an overall increase in response gain following salicylate (Noreña & Farley, 2013; Auerbach *et al*., 2014).

It was notable that the magnitude of the salicylate effect was significantly greater at the frequencies above and below the BF for that electrode site. This may reflect greater headroom for increased firing when units are driven less strongly by frequencies either side of BF. The changes in firing rate within the eFRA are consistent with either increased excitation or reduced inhibition. Importantly, we did not observe any systematic expansion of the eFRA in the presence of salicylate. Thus, our data suggest there is no reduction in the flanking inhibition that has been observed at frequencies above or below the eFRA for some neurons (Palombi & Caspary, 1996; Egorova *et al*., 2001; LeBeau *et al*., 2001; Palmer *et al*., 2013. Nevertheless, this interpretation is not definitive given that we recorded multi-rather than single units - a difference that may have obscured such changes in the FRA. Intriguingly, consistent with this finding, a recent fMRI study of tinnitus in humans reported a larger BOLD response to tones in tinnitus patients compared with controls to several frequencies above and below the BF for that location {Berlot, 2020 #6773).

Some previous studies have reported that systemic salicylate induces a threshold shift in the IC (Boettcher *et al*., 1989; Ma *et al*., 2006; Jiang *et al*., 2017). However, in keeping with Bancroft *et al*. (1991) and Sun *et al*. (2009), we found no consistent direction of change for estimates of CF, threshold, or bandwidth taken from the eFRAs following systemic salicylate.

### 4.2 Local perfusion of salicylate in the ICc does not mimic systemic salicylate

The mechanism and origin of tinnitus induced by systemic salicylate remains elusive. The primary effect of salicylate in inducing tinnitus is believed to be a reduction in cochlear output that elicits changes in the central auditory pathway (Oliver *et al*., 2001; Guitton *et al*., 2003; Greeson & Raphael, 2009) (Kakehata & Santos-Sacchi, 1996; Wu *et al*., 2010). However, salicylate may also exert direct effects on central structures, indeed there is some evidence that salicylate has direct effects in the IC (Wang *et al*., 2008; Hu *et al*., 2014; Patel & Zhang, 2014). Here we perfused a 100-fold range of salicylate concentrations to obtain levels in the ICc spanning those expected following the dose used systemically (Brien, 1993; Wu *et al*., 2013). This local perfusion did not replicate the effects of systemic administration. Thus, although there was an increase in spontaneous and sound-driven firing in the first hour of perfusion, the change was much smaller than the maximum increase following systemic salicylate. Moreover, extending the perfusion time and/or increasing concentration of the drug failed to increase the magnitude of this effect and indeed caused a marked and persistent decrease in spontaneous and sound-driven firing. We conclude that hyperactivity in the ICc following systemic salicylate requires its effects outside the IC, most likely in upstream auditory structures. In isolation, the direct effects of salicylate on IC neurons are not sufficient to induce increased firing rates acutely.

### 4.3 Blocking nitric oxide signalling in the IC prevents salicylate-evoked hyperactivity

Strikingly, the salicylate-induced increase in both spontaneous and sound-driven firing was completely prevented by inhibiting the synthesis of nitric oxide in the IC by the local perfusion of L-MeArg at a concentration known to inhibit nNOS (Garthwaite *et al*., 1989). Comparison of the FRAs following salicylate alone with those in which salicylate was accompanied by perfusion of the IC with L-MeArg showed that L-MeArg abolished the increased driven response at all frequencies and levels within the FRA demonstrating that its effects were not frequency selective.

It is important to note that animals in which systemic salicylate was administered alone also had a microdialysis probe implanted in the ICc that was perfused with aCSF throughout. Hence, the difference between responses in these animals and those in which L-MeArg was perfused into the ICc can only be ascribed to the presence/absence of L-MeArg and, by extension, to the inhibition of nNOS in the ICc. Thus, the effects of systemic salicylate on spontaneous and driven firing within the ICc depend on nitric oxide signalling in the ICc. Since virtually all nNOS in the ICc is structurally and functionally linked to NMDA receptors (Olthof et al., 2019), it is reasonable to assume that the nNOS-dependent response to salicylate involves activation of these nNOS-linked NMDA receptors. Indeed, the local perfusion of NMDA in the ICc evokes increases in spontaneous and sound-driven firing reminiscent of those elicited by systemic salicylate and they are similarly blocked by local perfusion of L-MeArg (Olthof *et al*., 2019). Thus, the nNOS-dependent response to salicylate is likely mediated by an increase in the activation of these nNOS-linked NMDA receptors presumably by increased activity in ascending glutamatergic inputs.

### 4.4 Mechanistic considerations

An important question is the source of the increased drive to these NMDA receptors. The nNOS linked NMDA receptors on neurons in the ICc are contacted by a specific population of glutamatergic terminals that contain both vesicular glutamate transporter 1 and 2 (VGLUT1 and VGLUT2) (Olthof *et al*., 2019). These likely originate from the T-stellate neurons in the ventral cochlear nucleus (Ito & Oliver, 2010; Ito *et al*., 2011). T-stellate cells are connected in networks and are proposed to generate gain control mediated by NO: a physiological mechanism that enables these cells to enhance and maintain spectral contrast over a wide range of levels (Cao *et al*., 2019). Acoustic trauma leads to the over excitability of T-stellate neurons (Cai *et al*., 2009) and, consistent with this, they show increased expression of nNOS in animals with tinnitus induced by noise exposure or salicylate (Zheng *et al*., 2006; Coomber *et al*., 2014; Coomber *et al*.). Increased membrane expression of both nNOS and the NR1 NMDA receptor subunit has been reported in the ventral cochlear nucleus following cochlear ablation (Chen *et al*., 2004). Interestingly, recent evidence also reveals an increased neuronal modulation by nitric oxide in the ventral cochlear nucleus in animals with behavioural evidence of tinnitus following noise exposure (Hockley *et al*., 2020a; Hockley *et al*., 2020b).

We speculate that the pathological plasticity induced by salicylate leads to increased firing in T-stellate inputs to the ICc, resulting in the overstimulation of nNOS-linked NMDA-receptors, and causing enhanced spontaneous firing and hyperactive responses to sound. Such changes may lead to pathological plasticity in the IC itself, since animals treated with salicylate show increased IC expression of the NMDA-receptor subunit NR2B (Hu *et al*., 2014). However, the finding that salicylate perfused locally in the IC did not result in hyperactivity suggests that its effects on the IC alone is insufficient to induce plasticity in the absence of changes elsewhere in the auditory pathway.

Although our findings suggest that the predominant effects of salicylate in the IC result from increased excitatory drive, other studies have demonstrated that systemic salicylate, and other manipulations likely to induce tinnitus, modulate other neurotransmitter mechanisms in the IC including GABAergic inhibition and serotonergic mechanisms (Bledsoe *et al*., 1995; Suneja *et al*., 1998; Bauer *et al*., 2000; Liu *et al*., 2003; Dong *et al*., 2010b; Patel & Zhang, 2014). In the case of systemic salicylate administration, the timescales of the effects may be important. Our results are based on acute exposure to salicylate over a few hours whereas changes in GABAergic measures were observed after exposure to salicylate over many weeks (Bauer *et al*., 2000). It is possible that modulation of inhibition is important in tinnitus and that the acute changes in excitatory drive observed here induce long-term changes in inhibitory mechanisms that are necessary to establish or maintain chronic tinnitus.

### 4.5 Implications for tinnitus in humans

An important consideration is the extent to which our hypotheses and observations using systemically administered salicylate are relevant to tinnitus and hyperacusis in humans that can have many causes. As we have discussed above, increased activity in higher auditory structures is a feature of tinnitus induced experimentally by salicylate, noise exposure, and mechanical lesions of the cochlea, suggesting it is a general feature of tinnitus rather than specific to salicylate (Chen *et al*., 2004; Coomber *et al*., 2015). The rapid development of high field strength fMRI with the spatial resolution to image small structures makes it increasingly possible to study human IC (De Martino *et al*., 2013; Berlot *et al*., 2020). Such methods will be useful in discovering how far findings in various animal models generalise to human tinnitus.

### 4.6 Conclusion

Based on our findings, we propose a network in the ascending auditory pathway that modulates gain control in the IC via NMDA receptors and nitric oxide that in pathology may underpin tinnitus. The components of this pathway in the ICc - NMDA-receptors, nNOS, the downstream effector soluble guanylyl cyclase, and the PSD-95 anchor protein may represent druggable targets for the treatment of tinnitus.

## Acknowledgments

BMJO was supported in part by a PhD studentship from Newcastle University. We thank Nick Lesica for generously making available his software for sound stimulation and electrophysiological recording and John Garthwaite for advice on drugs for modulating the nNOS pathway. We are grateful for financial support from the BBSRC (BB/J008680/1 to AR and BB/P003249/1 to AR and SEG), and Flexigrant F38 from Action on Hearing Loss to AR.

## Notes

***Conflict of interest*:** The authors declare no competing financial interests.

### Competing Interest Statement

The authors have declared no competing interest.

